# Three new species of *Thelymitra* (Diurideae, Orchidaceae) endemic to Aotearoa New Zealand

**DOI:** 10.64898/2026.08.27.745643

**Authors:** H.R. Jones, J.A. Tate, C.A. Lehnebach

## Abstract

Three new species of sun orchid (*Thelymitra*) endemic to Aotearoa New Zealand are here described: *T. palustris*, *T. scabrifolia* and *T. semaphora*. The morphological distinctiveness of these three species has been acknowledged for decades; however, their taxonomic status has remained unresolved. Evidence from existing karyological data, recently generated DNA sequence data (*LFY* and *ycf*1) and morphological studies from historical and fresh collections are used here to support their formal description. Both *T. palustris* and *T. semaphora* are restricted to wet habitats north of Auckland (North Island). *Thelymitra scabrifolia* inhabits mostly scrub, with a similar northern North Island distribution, but it has been found also in Manawatāwhi / Three Kings Islands and historically in Otago (South Island). All three species are polyploids and are of conservation concern.

## Introduction

The genus *Thelymitra* J.R.Forst. & G.Forst. (Diurideae; Orchidoideae) – otherwise known as sun orchids, is a diverse group of terrestrial orchids that originated in Australasia almost 10 Ma (Nauheimer et al. 2018). It includes more than 120 species (POWO, 2026), with most species occurring in Australia (> 100 species and eight hybrids) (AVH 2026) and Aotearoa New Zealand (NZ) (15 species, one hybrid) (Allan Herbarium 2025). Only one or two species have been recorded elsewhere, such as East Timor, Indonesia, New Caledonia, New Guinea, and the Philippines (Nauheimer et al. 2018).

Unlike many other orchid genera, in *Thelymitra* the labellum resembles the other sepals and petals in shape, size and colour, giving the flowers a radially symmetrical appearance (see Figure 1A, D, G). The column morphology, on the other hand, is more complex than the labellum in these species. A structure at the apex of the column, known as the mitra, is formed by the fusion of the staminodes, style, and anthers (Fig. 1B, E, H; Burns-Balogh and Bernhardt 1988). This structure is taxonomically informative and has been previously used to support species circumscription and assist with identification (Moore 1968; Jeanes 2004, Rolfe and de Lange 2010; Jones and Lehnebach 2022). Another unique feature of *Thelymitra* is the reliance on windless, warm and sunny conditions for their flowers to open, a feature that gave rise to their common name, sun orchid.

**Figure 1:**
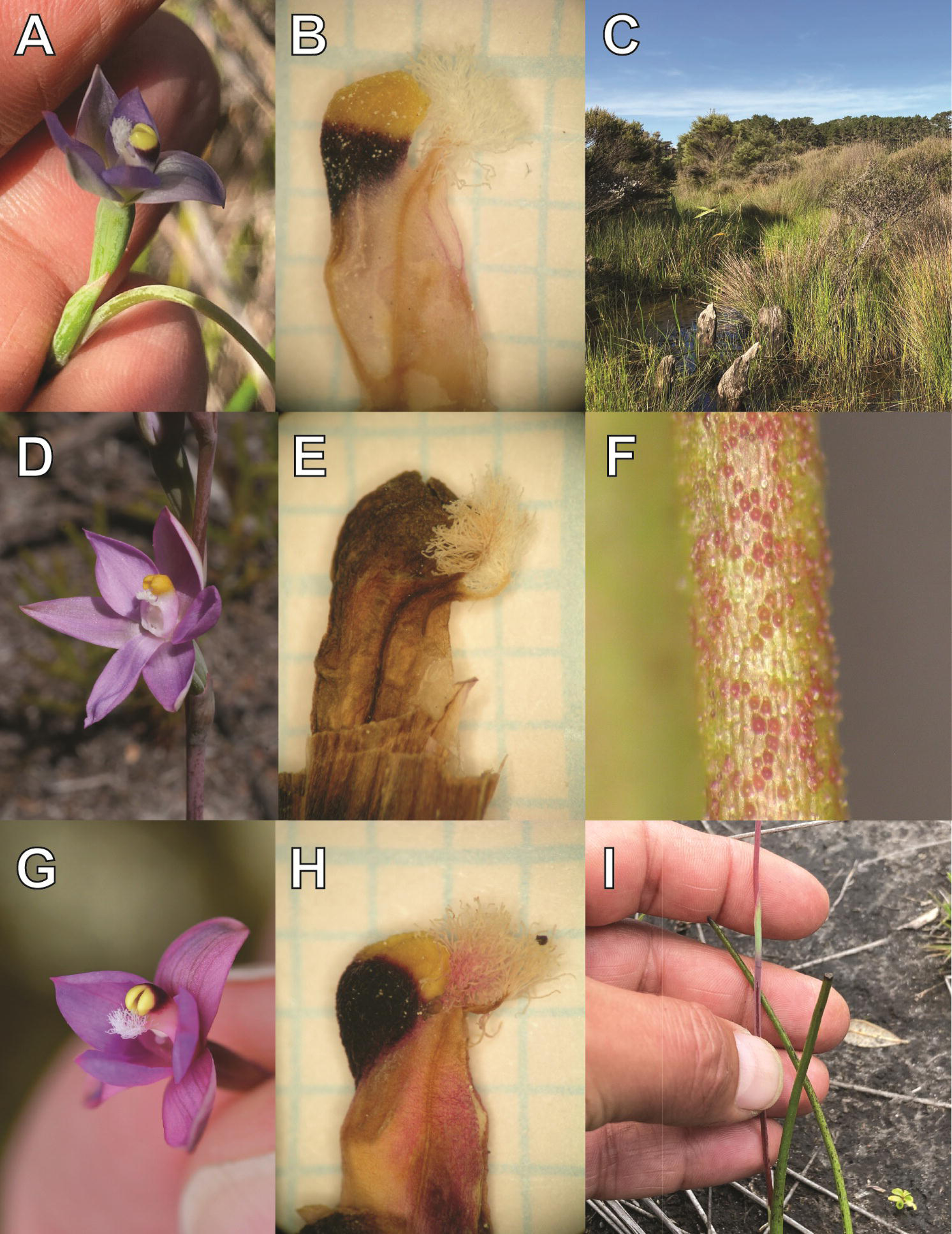
Three new species of sun orchid (*Thelymitra*) endemic to Aotearoa New Zealand. *Thelymitra palustris* (A: flower, B: column, C: habitat), *T. scabrifolia* (D: flower, E: column, F: leaf margin and underside) and *T. semaphora* (G: flower, H: column, I: red stem and bright green stem bract). Scale bar in A, D, G ∼1cm. Grid in paper graph in the background of B, E, F is 1 mm. B (WELT SP 111004), E (WELT SP119844), F (WELT SP121775), H (CHR306812).

Phylogenetic and biogeographic studies by Nauheimer et al. (2018) suggest long-distance dispersal from Australia, assisted by the prevailing westerly winds, was involved in the arrival of *Thelymitra* in NZ. Their study also showed that hybridisation played an important role in the diversification of this genus. In the past, hybridisation and polyploidy have been linked to the evolution of some NZ *Thelymitra* (Molloy and Dawson 1998, Peakall and Molloy 1998, Dawson et al. 2007), but only recently this was confirmed by Jones et al. (2025) who produced the most complete phylogenetic study of NZ *Thelymitra*. Their work uncovered an extensive history of polyploidy and hybrid speciation across NZ sun orchids and confirmed the allopolyploid nature of one native and three endemic species. These are *T. carnea* (4*n* = 62, *T*. *flexuosa* × *T. pauciflora*), *T. hatchii* (4*n* = 66, *T. formosa* × *T. longifolia*), *T. pulchella* (4*n* = 66, *T. cyanea* × *T. longifolia*) and *T. nervosa* (4*n* = 54, *T*. *ixioides* × *T*. *longifolia*).

In addition, Jones et al. (2025) showed that three taxonomically unresolved entities with high chromosomes counts (i.e. 2*n* = 60 & 2*n* = 84) that are currently known in NZ by tag-names only are likely of allopolyploid origin. These taxa are *Thelymitra* “Ahipara” (WELT SP079140), *T*. “darkie” (CHR 518036), and *T*. “rough leaf” (AK 22953). Despite the morphological distinctiveness of these entities having been recognised for decades, and the general agreement among New Zealand botanists that they may represent valid species (Rolfe and de Lange 2010), they have remained undescribed. Cytological evidence from earlier studies (Hair 1942, Dawson et al. 2007) similarly supported their distinctiveness.

Here we describe three new species of *Thelymitra* endemic to Aotearoa NZ supported by a combination of historical specimens, newly collected morphological data and the genetic evidence generated by Jones et al. (2025). Currently, all three species are of conservation concern (de Lange et al. 2024), but due to their unresolved taxonomic status they have not been included in formal conservation programmes. By resolving their taxonomic status and providing formal descriptions this work will assist the Department of Conservation in undertaking targeted demographic surveys, managing populations effectively, and guiding future conservation initiatives.

## Methods

### Morphological study

We studied fresh and historical specimens of *Thelymitra* at AK, CHR, MPN, and WELT herbaria. Herbarium acronyms follow Thiers (2026). We examined specimens of *Thelymitra* “Ahipara” (N=40), *Thelymitra* “darkie” (N=13), and *Thelymitra* “rough leaf” (N=16) using an Olympus stereomicroscope and measured them using a digital calliper. Morphological characterisation for each species was further supplemented by observations available on iNaturalist (https://inaturalist.nz/).

Following Moore (1968, 1970), Molloy and Hatch (1990), Jeanes (2004), and Rolfe and de Lange (2010), we studied characters considered taxonomically informative, such as leaf dimensions and surface, stem morphology and number and colour of stem bracts, and morphology and colour of the post-anther lobe. Overall, we focused on 14 characters, including vegetative and floral characters.

Flowering times and geographic distribution for each species were determined from herbarium records, field observations and observations reported on iNaturalist (https://inaturalist.nz/). In total we included 52 observations from iNaturalist reported by 13/08/2025 (Supplementary Table 1).

### Phylogenetic affinities

Phylogenetic affinities of all three entities were previously assessed and discussed *in extenso* by Jones et al. (2025), which was based on sequences from the chloroplast region *ycf*1 and the single-copy nuclear gene *LFY*. We used these sequences and a selection of the taxa used by Jones et al. (2025) to construct a phylogenetic network for each data set using the Neighbour Net algorithm in SplitsTree 6.0.0 (Huson and Bryant 2024). The network was constructed with constant sites omitted and uncorrected p-distances. These networks (Figure 2A & B) are intended only to provide a visual framework for the phylogenetic affinities of each new species as relationships among Australian and NZ *Thelymitra* were discussed previously in Jones et al (2025).

**Figure 2:**
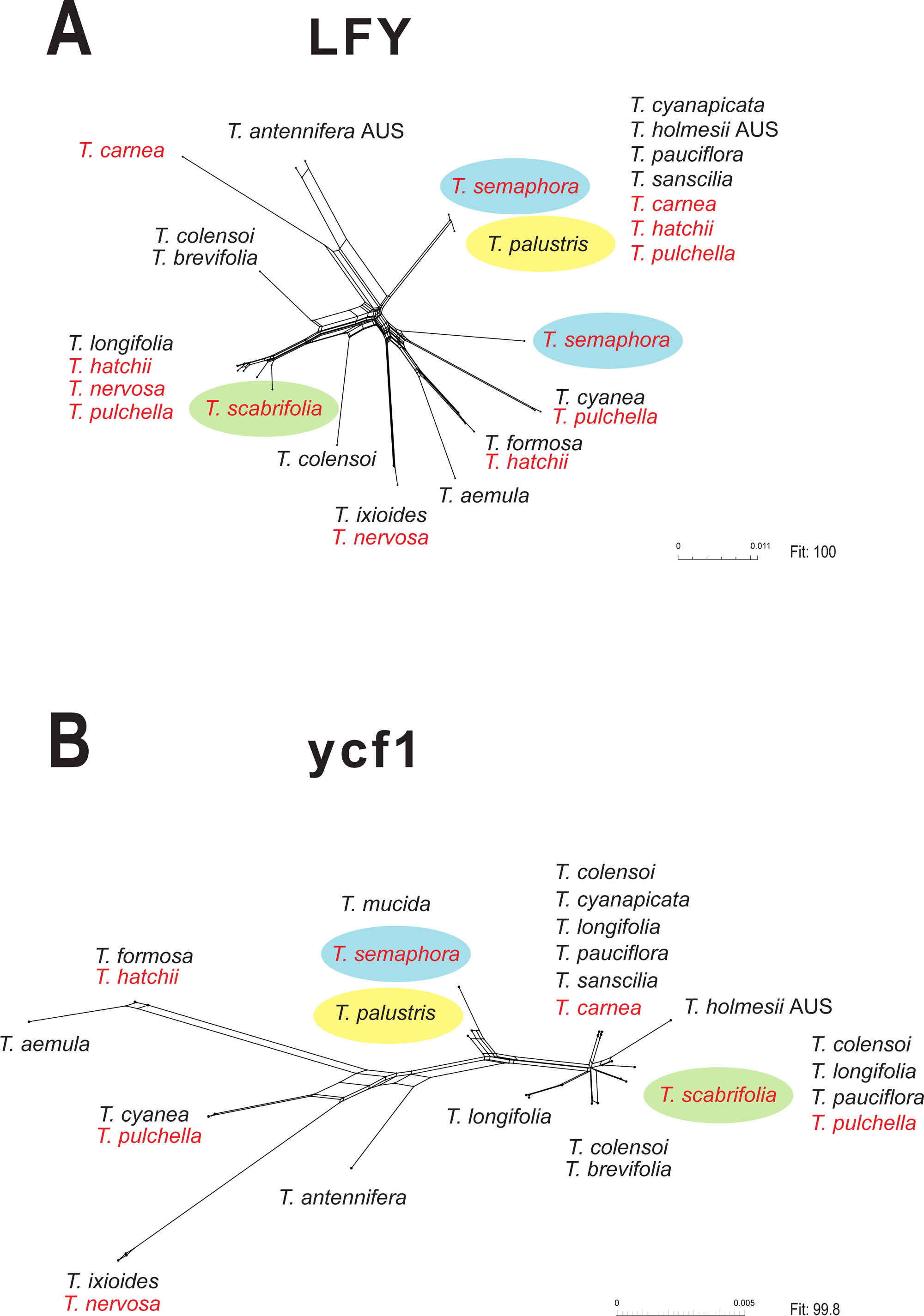
Phylogenetic affinities of *Thelymitra palustris*, *T. scabrifolia* and *T. semaphora* based on nuclear (*LFY*) (A) and plastid (*ycf1*) (B) sequence data generated by Jones et al. (2025). New species are indicated with coloured ellipsoids. Polyploid species are indicated in red font. AUS: Australia.

## Results

### Taxonomy

*Thelymitra palustris* H.R.Jones & Lehnebach, *sp. nov.* (Figure 1A – B)

**Common name**: Ahipara swamp sun orchid

**Type**: NORTH ISLAND: Northland, Ahipara. Sandhills Road. 15 November 1990, *de Lange P. 528* & *Crowcroft G*. *s*.*n*. (holotype: WELT SP079140!). Figure 3

**Figure 3:**
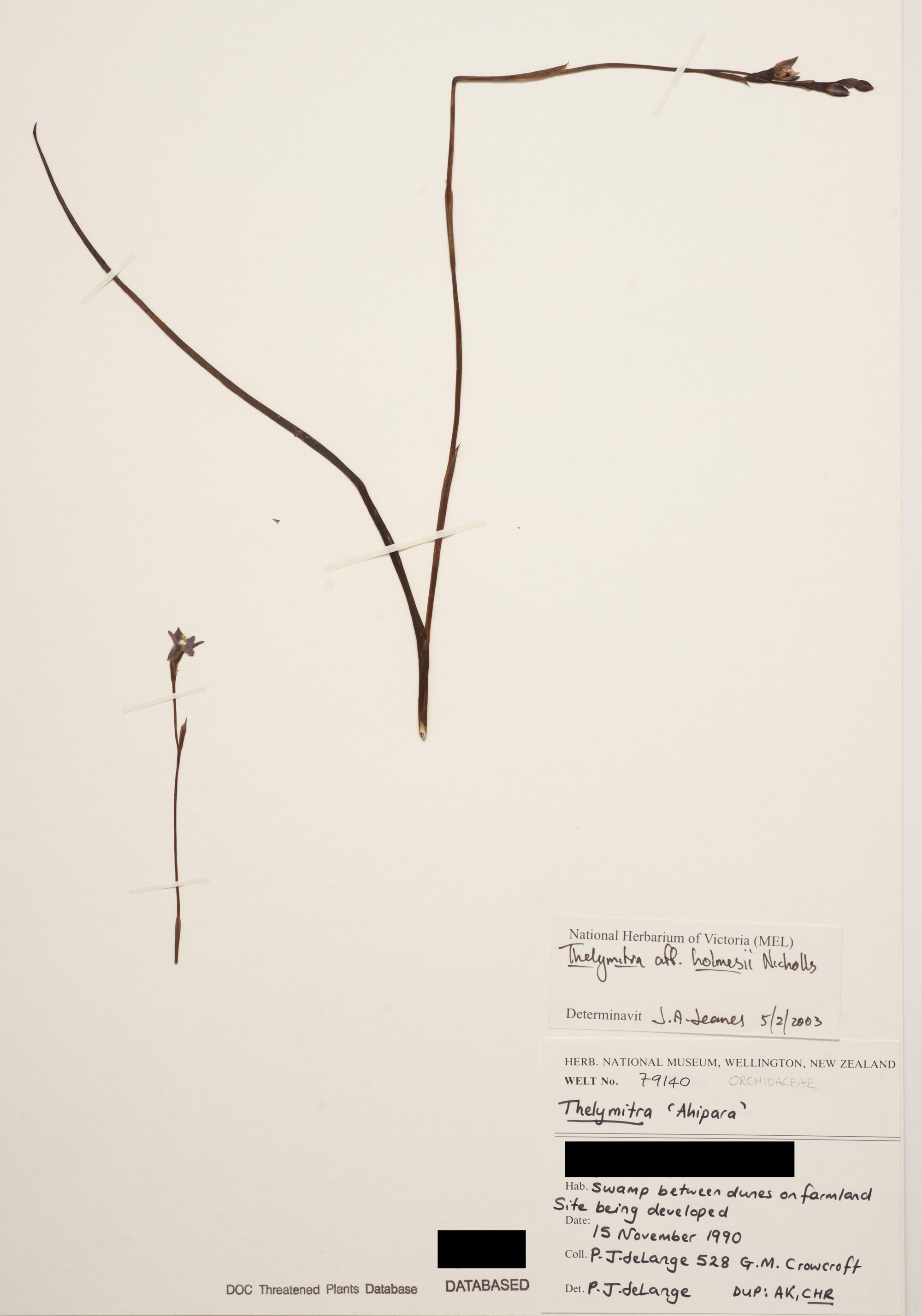
Holotype of *Thelymitra palustris* H.R.Jones & Lehnebach at WELT Herbarium, Museum of New Zealand Te Papa Tongarewa. Accession number WELT SP079140.

**Iconography**: Tyler and St. George (2007, pp. 724– 729).

**Diagnosis**: *Thelymitra palustris* resembles some forms of *T. pauciflora* in its blue flowers and split post-anther lobe but it differs in the orientation and coloration of the latter. In *T. palustris*, the post-anther lobe is predominantly ascending (vs dorsally compressed in *T. pauciflora*), topped with a yellow band that extends from below the point of bifurcation to the apex of each sub-lobe (vs originating above the point of bifurcation in *T. pauciflora*).

#### Description

**Plant** 20 – 64 cm tall at flowering, shorter when growing in exposed habitat. **Leaf** (60 –) 100 – 300 (–350) × (1.8–) 3 – 7 (–9) mm, solitary, erect or lax, linear-lanceolate, light green with a reddish base, leaf blade strongly V-shaped in the lower half and then slightly flattened toward the leaf apex, apex acute. **Stem** (1–) 1.6 – 3.6 mm diameter, slender to moderately robust, glaucescent to pale purple. **Basal sheath** up to 35 mm long, pale, truncate, mucronate. **Stem bracts** usually 2 – 3, (15.5 –) 18 – 60 (– 100) × 2 – 5 (– 6.8) mm, bright green, lanceolate, apex acuminate. **Floral bracts** 6 – 16 mm long, lanceolate, apex acuminate-aristate. **Flowers** (1–) 2 – 6 (–7), 10 – 15 mm diameter, pale blue, held on slender 7.5 – 12.3 mm long pedicels, partially covered by a bract. **Ovary** 6.9 – 8.7 × 3.0 – 3.5 mm, oblanceolate, terete. **Perianth segments** concave, ovate, acute to shortly apiculate. **Sepals** 9.0 – 10.5 ×4.0 – 5.8 mm (dorsal) and 9.4 – 10.7 × 3.5 – 5.1 mm (lateral). External and internal surfaces pale green to green towards the center, margins mostly white towards the base grading to pale blue towards the reddish apex. **Petals** 8.0 – 10.6 × 4 – 5.5 mm (lateral) and 8.5 – 9.9 × 3.8 – 5.9 (labellum). External and internal surfaces blue to pale blue - white towards the base and red-pink at the apex. **Column** up to 5 mm tall, erect, concave in side view, white to pale mauve. **Post-anther lobe** as high or exceeding the anther, horseshoe shaped, split into two sub-lobes, dark reddish-brown below the point of bifurcation and bright yellow above this point towards the apex of each sub-lobe. **Column arms** terete, curved upwards, adorned at the distal end with white cilia almost as long as the arm. **Capsule** 10.5 – 13.5 × 3.8 – 5.8 mm, obovoid. **Pollen**: monads.

**Etymology**: The epithet is derived from the Latin word *palus* which means swamp/marsh and it refers to the wet habitat where this orchid generally grows.

**Distribution**: Endemic to New Zealand. Restricted to the Aupouri Ecological District of the Far North in the North Island. Historically found near Ahipara township, on the west coast of the North Island. Extant populations near Kaimaumau and Waipapakauri Beach.

**Habitat**: In wetlands (Figure 1C). Growing in the ecotone between sand dunes and peaty ponds. On damp swampy ground or rotting *Agathis australis* (D.Don) Lindl. stumps slightly above the water level. Some of the dominant vegetation includes *Empodisma robustum* Wagstaff et B.R. Clarkson (Restionaceae), *Gleichenia microphylla* R.Br. (Gleicheniaceae), *Leptospermum hoipolloi* f. *incanum* (Cockayne) de Lange & L.M.H Schmid (Myrtaceae), *Machaerina teretifolia* (R.Br.) Koyama (Cyperaceae), and *Schoenus brevifolius* R.Br (Cyperaceae).

**Flowering**: October – November

**Chromosome number**: 2*n* = 60. (Dawson et al. 2007). Voucher for chromosome count: CHR584289.

**Notes**: The first record of *Thelymitra palustris* dates from 1987 when Doug McCrae (1896 – 1990) and Brian Molloy (1930 – 1922) found it in wetlands near Ahipara township (McCrae 1987, de Lange et al. 1991). From then onwards this species was known as *T*. “Ahipara”. At the time, and based on available evidence, *T*. “Ahipara” was treated as a possible NZ endemic (de Lange et al 1991) and most currently it has been associated with the *T. pauciflora* aggregate (Rolfe & de Lange 2010). de Lange et al. (1991) also highlighted the similarity between *T*. *palustris* and *T. “*darkie*”*, another undescribed species that we formally describe below. Decades later the affinity between these two entities was confirmed by cytological data (Dawson et al. 2007) and chloroplast and nuclear DNA sequences (Jones et al. 2025). The latter study, which also included Australian representatives of *Thelymitra*, revealed that samples of two Australian endemics *T*. aff. *holmesii* Nicholls and *T. mucida* Fitzg., were also phylogenetically close to *T. palustris* in the *LFY* and *ycf*1 phylogeny, respectively (Figure 2A, B). Similar to *T. palustris*, both Australian species prefer wet-swampy soils.

In the past *T. palustris* has been likened with *T. holmesii* and Australian botanist Jeffrey A. Jeanes has acknowledged the morphological similarity between these two species (see determinative label on WELT SP079140, Figure 3), however they differ in chromosome numbers (i.e. *T. palustris* 2*n*=60 vs *T. holmesii* 2*n*=57, Dawson et al. 2007) and *ycf*1 sequences (Jones et al. 2025).

An allopolyploid origin for *T. palustris* is likely. The study by Jones et al. (2025) revealed discordance in the position of *T. palustris* between nuclear and chloroplast phylogenies (see also Figure 2A, B). A similar origin has been reported and confirmed for other NZ *Thelymitra* (Molloy & Dawson 1998, Jones et al 2025), but unlike those, the putative progenitors of *T. palustris* have not been yet found in NZ. Considering its close affinity with the Australian species *T. mucida* and *T. holmesii*, together with records of putative hybrids involving *T. holmesii* in Australia (Weber & Entwisle 1994), it is plausible that *T. palustris* originated in Australia and subsequently dispersed to NZ. This scenario was hypothesised for *T. palustris* when its chromosome numbers were first reported by Molloy & Dawson (1998). A similar process was proposed for the evolution of *T. carnea*, which is an allopolyploid now established in NZ that originated in Australia after hybridisation of *T. flexuosa* and *T. pauciflora* (Molloy & Dawson 1998).

**Conservation status**: Threatened – Nationally Critical (de Lange et al. 2024). This status means the species is represented by a very small population, due to natural or unnatural causes, and that the total area of occupancy is ≤ 1ha (0.01 km^2^). The main threat for the survival of this species is habitat destruction, i.e. wetland drainage. The effect of wildfires, as those occurring in 2021 in Kaimaumau (RNZ 2021), on adult plants and recruitment is unknown.

*Thelymitra palustris* is one of the few examples of orchid translocation that have taken place in NZ. This action was prompted by the imminent impact habitat transformation (i.e. conversion of wetlands into pasture) would have had on the only known population. To prevent complete extermination of this taxon, nearly 400 plants were transferred from its original locality to suitable wetlands near Lake Ohia and the Ahipara Gumfields (de Lange et al 1991). To the best of our knowledge, it seems the fate of these plants was not followed up as no public document reporting survival records exists.

Propagation using symbiotic or asymbiotic methods could assist with the conservation of *T. palustris*, but first its mycorrhizal associations and pollination system should be investigated.

**Selected specimens studied**: NORTH ISLAND: Aupouri, [near] Ahipara, *de Lange P*. *J*. and

*Crowfort G. M*. *s*.*n*., 15 Nov 1990, AK 200883, AK 200884, WELT SP079140. Aupouri, *ex*. Sand Hills Rd [Cultivated at Auckland University], *Cameron E*. *K*. *s*.*n*. 19 Nov 1990, AK 234660. Aupouri, Karikari Peninsula, Lake Rotokawau, *de Lange P. J. 519 & Crowcroft G. M.*, 16 Nov 1990, CHR 47303. Aupouri, [near] Kaimaumau, *Hooper K*., *Campbell B*. & *Townsend A*. *s*.*n*., 2 Dec 2020, WELT SP110814. Aupouri, [near] Kaimaumau, *Hooper G*., *Campbell B*. & *Townsend A*. *s*.*n*., 2 Dec 2020, WELT SP110816, Aupouri, near Waipapakauri Beach, *Lehnebach C*.*A*., *Zeller*, *A*.*J*. and *Matthews K*., WELT SP 111004.

Selected observations reported on iNaturalist:

http://naturewatch.org.nz/observations/4620619,

https://www.inaturalist.org/observations/36649238,

https://www.inaturalist.org/observations/36649894,

https://www.inaturalist.org/observations/66367266.

*Thelymitra scabrifolia* H.R.Jones & Lehnebach, *sp. nov.* (Figure 1 D – F)

**Common name**: Rough leaf sun orchid

**Type**: NORTH ISLAND: Northland, Motutangi Swamp, *McCrae, D. P. s*. *n*. [Cultivated at Taylor Rd, near Paranui, North Auckland and Lincoln, Landcare Research], (holotype: CHR 645411!) Figure 4

**Figure 4:**
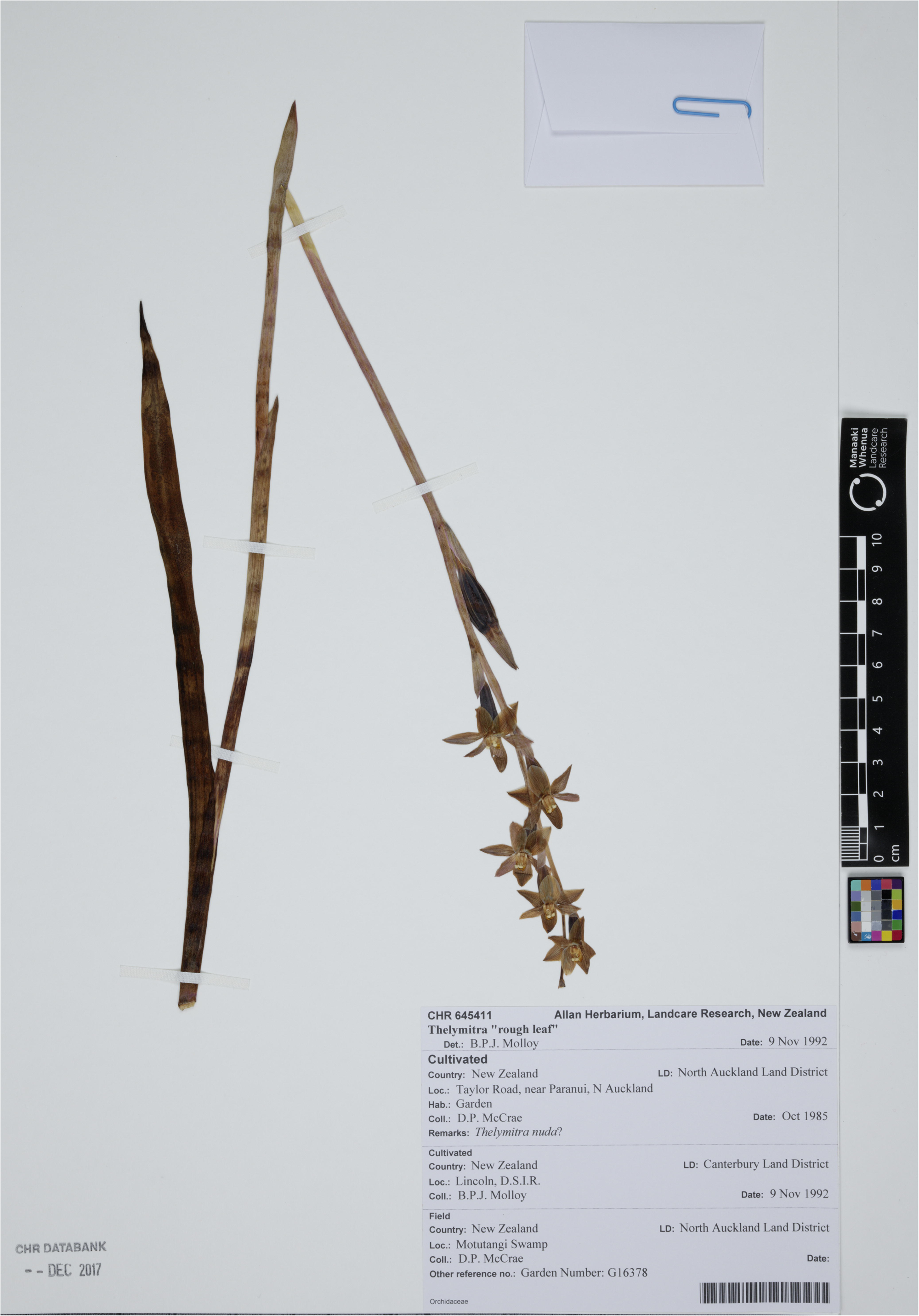
Holotype of *Thelymitra scabrifolia* H.R.Jones & Lehnebach at the Allan Herbarium (CHR). Accession number CHR 645411. © New Zealand Institute for Bioeconomy Science Limited, 2026. CC BY 4.0.

**Iconography**: St George et al. (1996, p. 121), Tyler and St. George (2007, pp. 741– 747).

**Diagnosis**: *Thelymitra scabrifolia* resembles some forms of *T. longifolia* in having column arms bearing dense, plumose cilia and its almost entire, cucullate, yellow-orange post-anther lobe, but differs in having mauve to pale pink flowers. *Thelymitra scabrifolia* differs from all NZ *Thelymitra* by its rough abaxial leaf surface.

#### Description

**Plant** 22 – 52 cm tall at flowering, shorter when growing in exposed habitat. **Leaf** (180 –) 210 – 320 (– 380) × 4.6 – 10 (–14.3) mm, solitary, erect or lax, linear-lanceolate, yellow-green, lamina channelled with abaxial surface tuberculate, apex acute. **Stem** (1.5 –) 2.3 – 4.1 mm diameter, slender to moderately robust, reddish-brown. **Basal sheath** up to 35 mm long, pale, truncate, mucronate. **Stem bracts** usually 1 – 2, 33 – 47 (– 65) × (3.8 –) 5 – 7.4 (– 9.0) mm, clasping the stem, bright green at the base and reddish brown towards the lanceolate-acuminate apex. **Floral bracts** 10 – 20 mm long, green or reddish brown, lanceolate, apex acuminate-aristate. **Flowers** (2 –) 4 – 8 (– 12), 20 – 40 mm diameter, held on slender 4.9 – 9.4 mm long pedicels, partially covered by a bract. **Ovary** 10.4 – 18.6 × 3.7– 7 mm, oblanceolate, terete. **Perianth segments** concave, ovate, acute to shortly apiculate. **Sepals** 9 – 18.5 × 4.0 – 7.0 mm (dorsal) and 12 – 17.5 × 5.5 – 6.0 mm (lateral). External surface reddish-brown or dull green towards the center, margins mostly pale pink or white. Internal surface pale pink or mauve. **Petals** 11 – 14.7 × 6 – 6.9 mm (lateral) and 8.1 – 15.4 × 5.3 – 6.1 (labellum). External and internal surfaces pink to pale mauve or pale blue. **Column** up to 4.7 mm tall, erect, concave in side view, white to pale pink. **Post-anther lobe** cucullate, bright yellow with brown margin, apex entire to slightly emarginate. **Column arms** terete, curved upwards, adorned at the distal end with white firm cilia. **Capsule** 15 – 20 × 6 – 8 mm, obovoid. **Pollen**: monads.

**Etymology**: The epithet is derived from the Latin words *scaber* (rough) and *folium* (leaf). The epithet refers to the rough texture of the leaf blade and aligns with the tag-name currently used for this orchid.

**Distribution**: Endemic to New Zealand. Mostly found in the North Island, from Te Paki to Eastern and Western Northland ecological regions. Also present on Manawatāwhi / Three Kings Islands. Historical collections suggest it is also found in the South Island (Otago).

**Habitat**: Found from coastal to lowland habitats. Usually in open shrubland, on clay pans, gumland scrub, forest margins and ultramafic shrubland. Mostly associated with rotting kauri wood, kauri podzols (highly acidic, nutrient-poor soils) and exposed ferricrete pans (iron-rich layers of soil or rock). Also found on bare ground under *Kunzea ericoides* (A.Rich.) Joy Thomps (Myrtaceae).

**Flowering**: October

**Chromosome Numbers**: 2*n* = 84 (Dawson et al. 2007). Vouchers for chromosome counts: CHR 584338, CHR 584339.

**Conservation Status**: At Risk – Naturally uncommon (de Lange et al. 2024). This category includes taxa whose distribution is naturally (i.e. not due to human disturbance) confined to specific substrates, habitats or geographic areas or that occur within natural small and widely distributed populations (Townsend et al. 2008). Although threats to the conservation of this species have not been identified, it is likely habitat destruction has adversely affected its distribution and abundance, as many of the recent records are from areas with less than 30% of indigenous biodiversity remaining and severely fragmented habitats (Cieraad et al. 2015).

**Notes**: The earliest published record of *Thelymitra scabrifolia* we found dates from 1987. This occurs in a list compiled by Doug McCrae (1896 – 1990) where *T*. “rough leaf” is mentioned along other orchids found at Kaimaumau wetland by members of the Kaitaia Orchid Society (McCare 1987). Since then, *T. scabrifolia* has been considered unlike any other species of *Thelymitra* in NZ and Rolfe & de Lange (2010) have hypothesised it has no close affinities with any other NZ species.

The most recent phylogenetic assessment of NZ *Thelymitra* by Jones et al. (2025) did not fully resolve the origin of *T. scabrifolia*, but revealed affinities with *T*. *colensoi*, *T*. *longifolia*, *T*. *pauciflora*, the allopolyploid *T. pulchella*, (Figure 2A, B) and Australian members of the *T. pauciflora* aggregate such as *T. albiflora* and *T. arenaria*. Its affinity with *T. longifolia* is not unexpected given the superficial similarity of *T. scabrifolia*’s flowers (e.g. petal colour and post anther lobe shape and colour) to those of some forms of *T. longifolia*.

Until recently, the rough leaf texture of *T. scabrifolia* (Figure 1 F) had not been reported for any other *Thelymitra*. A recently described Australian species, however, shares a similar feature (Mitchell 2026). The leaf of *Thelymitra asperifolia*, a member of the *T. pauciflora* aggregate, has been described as similar to an “*emery board*” (Mitchell 2026). It is not specified, however, whether this texture is on the upper or lower leaf surface. Unlike *T*. *scabrifolia*, the Australian species has a “*reddish-brown scape*”, “*red to maroon bracts*” and “*light purplish to purple with bluish streaks*” flowers with “*emarginate to deeply V-notched/bifid*” post-anther lobe (Mitchell 2026).

At 2*n* = 84, *T. scabrifolia* has one of the highest chromosome numbers recorded for *Thelymitra* in NZ (Dawson et al. 2007). The Tasmanian species *T. viridis* has the same chromosome number (Dawson et al. 2007). Molloy and Dawson (1998) hypothesised that *T. scabrifolia*, and the other two species formally described in this article, evolved in Australia and then dispersed to NZ, where they colonised narrower and more specific habitats. Past hybridisation and polyploidisation are the most likely processes underlying the origin for *T. scabrifolia*. Nauheimer et al. (2018) and Jones et al. (2025) presented evidence that supports these two processes as well, with discordant phylogenies and evidence of duplication in loci typically retained as single copy.

#### Selected specimens studied

THREE KINGS. Three Kings, Manawatāwhi (Great Island), above North West Bay. *de Lange P*. *J*. *3345*, 05 December 1996. AK 232966, CHR 490843. Three Kings, Manawatāwhi, above North West Bay. *McKenzie D. s. n*., 02 December 1995. AK 224941.

NORTH ISLAND. Te Paki, North Cape Scientific Reserve, Plateau, *de Lange P*. *J*. *3139*. 11 Oct 1996. AK 229531, CHR 487583. Te Paki, North Cape Scientific reserve, North Cape Plateau, *de Lange P*. *J*. *8701*, 19 Oct 2009, AK309824. Eastern Northland, Taurikura Bay, south slopes of mount Manaia, *Forester L*. *s*. *n*., 29 Sep 1997, AK 245994. Eastern Northland and Islands, black rocks off Moturoa Island, south-western Crater Rim Island, *Wright A*. *E*. 30 Oct 1990. AK231738. Aupouri, *ex*. Motutangi swamp, *McCrae D*. *P*. *s*. *n*. [Cultivated at Taylor Rd, near Paranui, North Auckland and Lincoln, Landcare Research], CHR 645411. *ex* Motutangi, *McCrae D*. *P*. *s*. *n*. [Cultivated at Taylor Rd, near Paranui, North Auckland and Lincoln, Landcare Research], CHR 688345. Aupouri, Kaimaumau Wetland, *Campbell B*. *s*. *n*., 16 Oct 2021, WELT SP119844, WELT SP119845, WELT SP119846. Northland, Puketi Forest Headquarters, *Riddell K*. *s*. *n*., 01 Nov 1999, CHR 645280.

SOUTH ISLAND. Rakanui, *ex* Shag Point, Palmerston, *St. George I. M*. *s. n.*, Nov 1987, [Cultivated at Lincoln, Landcare Research], CHR 584338 B, CHR 584339 C [CHR 584339 A in spirit collection not seen].

#### Selected observations reported on iNaturalist

https://www.inaturalist.org/observations/8538079,

https://www.inaturalist.org/observations/8538082,

https://www.inaturalist.org/observations/17358388,

https://www.inaturalist.org/observations/17372835,

https://www.inaturalist.org/observations/17761905,

https://www.inaturalist.org/observations/17762010,

https://www.inaturalist.org/observations/98452506,

https://www.inaturalist.org/observations/186446679,

https://www.inaturalist.org/observations/247079195, https://www.inaturalist.org/observations/247081787.

*Thelymitra semaphora* H.R.Jones & Lehnebach, *sp. nov.* (Figure 1 G – I)

**Common name**: Traffic light sun orchid

**Type**: NORTH ISLAND: Northland, Sweetwater Rd., Kaitaia. 03 November 1988, *Molloy, B*. *P. J. s*.*n*. (holotype: CHR 636773!, arrow) Figure 5

**Figure 5:**
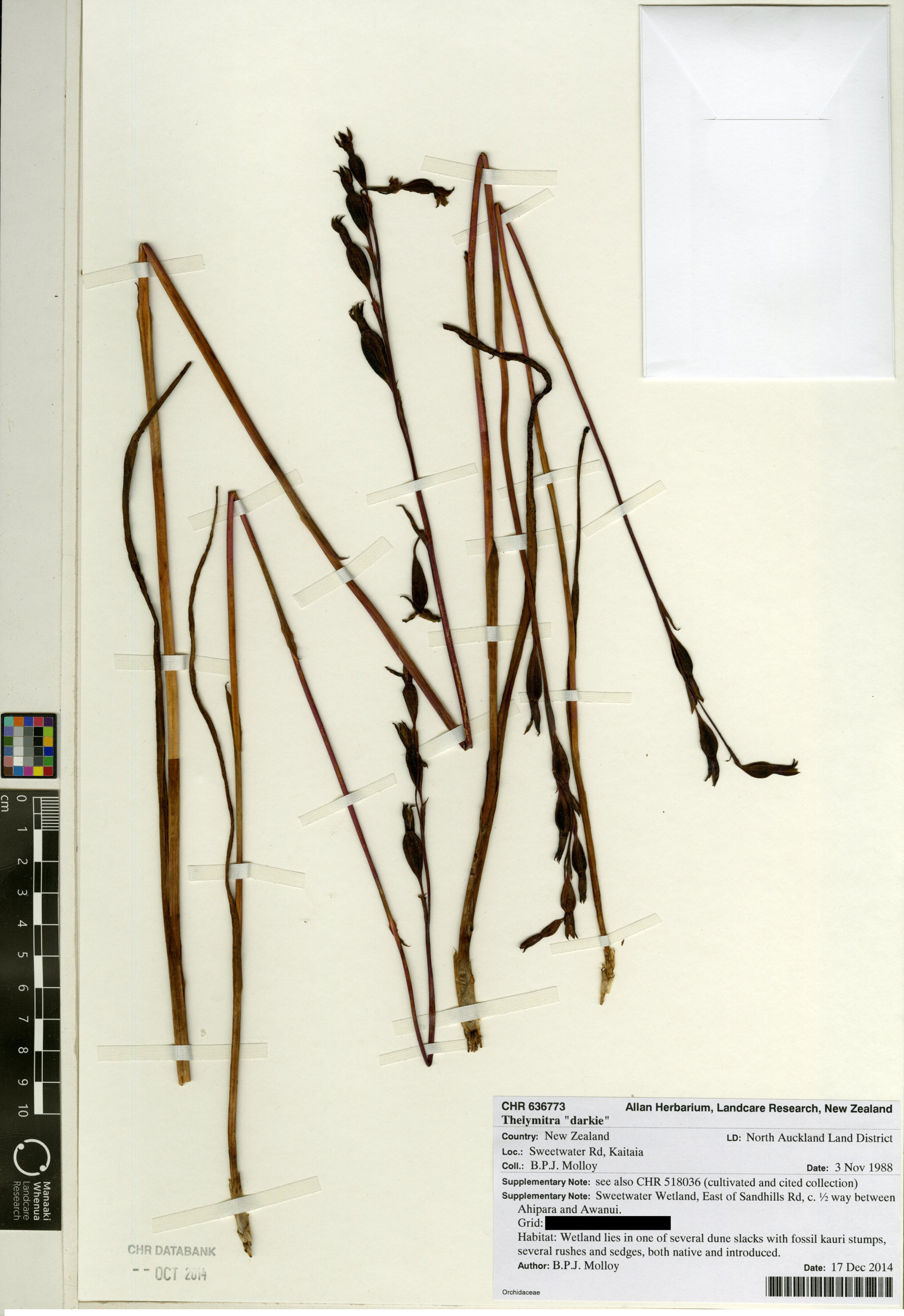
Holotype of *Thelymitra semaphora* H.R.Jones & Lehnebach at the Allan Herbarium (CHR). Accession number CHR 636773. © New Zealand Institute for Bioeconomy Science Limited, 2026. CC BY 4.0

**Iconography**: St George et al. (1996, p. 120), Tyler and St. George (2007, pp. 733– 740).

**Diagnosis**: *Thelymitra semaphora* most closely resembles *T. palustris* and *T. pauciflora* in the colour of its flowers but differs from both, and other known New Zealand *Thelymitra*, by its combination of reddish-brown stem and bright green stem bracts.

#### Description

**Plant** 20 – 66 cm tall at flowering. **Leaf** (90 –) 160 – 250 (– 440) × (2.4 –) 3 – 5.5 (– 8) mm, solitary, erect or spreading, linear-lanceolate, light green with a reddish base, leaf blade V-shaped in the lower half and then slightly flattened toward the leaf apex, apex acute. **Stem** (2 –) 3 – 4 mm diameter, moderately robust, reddish to brown-purple. **Basal sheath** 15 – 30 mm long, pale, truncate, mucronate or apiculate. **Stem bracts** usually 2 – 3, 23 – 50 (– 60) × (2 –) 3 – 5.4 (– 6.7) mm, bright green, lanceolate, apex acuminate. **Floral bracts** 8 – 14 mm long, lanceolate, apex acuminate. **Flowers** (1 –) 2 – 5 (– 7), 10 – 20 mm diameter, held on slender (8.5 –) 12 – 18 mm long pedicels, covered entirely by a bract. **Ovary** 10 – 19 × 3.6 – 5.3 (– 8) mm, oblanceolate, terete. **Perianth segments** concave, ovate, acute to shortly apiculate. **Sepals** 8.1 – 11 ×3.0 – 5.0 mm (dorsal) and 8.1 – 11 × 3.0 – 5.0 mm (lateral). External surface pale mauve to purple towards the center, margins mostly pink or white. Internal surface grading from pale pink to mauve or blue towards apex. **Petals** 7.7 – 10 × 3.1 – 6.0 mm (lateral) and 5.8 – 9.5 × 2.8 – 4.0 (labellum). External and internal surfaces bright pink, mauve or blue and red-pink at the apex. **Column** 4 – 6 mm tall, erect, concave in side view, pale mauve. **Post-anther lobe** exceeding the anther, horseshoe shaped, apex blunt not tapered, dark crimson – brown to almost black with bright yellow margin. **Column arms** terete, curved upwards, adorned at the distal end with thin mauve-white cilia. **Capsule** 13.0 – 18.0 × 5 – 8 mm, obovoid. **Pollen**: monads

**Etymology**: The epithet is derived from Greek words σ□μα (*s*ē*ma*) meaning sign, mark, signal, and φέρειν (*pherein*) meaning to carry, bear. It literally means “signal-bearing”. This is a reference to the reddish stem and bright green bracts along the stem which in combination resembles traffic light signals (i.e. semaphores).

**Distribution**: Endemic to New Zealand. Restricted to the North Island from Te Paki to the Auckland Ecological regions.

**Habitat**: Found in lowland scrub and wetlands habitats, especially gumlands, open ground, along wetland margins and rarely along tracksides in kauri (*Agathis australis*) forest.

**Flowering**: September - October

**Chromosome numbers**: 2*n* = 60 (Dawson et al. 2007). Voucher for chromosome counts: CHR 584261

**Conservation status**: At Risk – Declining (de Lange et al. 2024). Species in this category are not of major conservation concern as changes are normally buffered by a large total population size and/or a slow decline rate (see Townsend et al. 2008). However, the conservation status of *T. semaphora* has worsened since it was last assessed six years ago (see de Lange et al. 2018), when it was ranked as Not Threatened. If current decline continues, a change to list it under the Threatened category is likely. The causes for decline have not been documented, but habitat destruction are the most likely drivers.

Future conservation efforts should focus on determining total population size for this species and occurrence within conservation land. Understanding its mycorrhizal associations and dependency on pollinators for fruit set are two key research priorities that could help with translocations or reinforcement of declining populations.

**Notes**: *Thelymitra semaphora* was first reported in 1987 when Doug McCrae (1896 – 1990) and members of the Kaitaia Orchid society spent time surveying orchids at the Ahipara Gumfields Historic Reserve (McCrae 1987). It was noted that this “*new orchid*” had “*a number of features uncharacteristic of T. pauciflora*” (i.e. overall dark purple pigmentation of the stem and ovary, a feature that likely prompted the tag-name “darkie”) and an affinity for wet areas (McCrae 1987), not typically occupied by *T. pauciflora*. But, similarly to *T. palustris*, *T. semaphora* has long been included in the *T. pauciflora* aggregate (McCrae 1987, St. George et al. 1996, Rolfe & de Lange 2010).

In a personal communication cited by de Lange et al (1991), Brian Molloy considered *T. semaphora* to be related to *T. palustris* based on their shared chromosome number and breeding system. Close affinity between these two species has been further confirmed by phylogenetic studies using nuclear and chloroplast sequences (Jones et al 2025). Jones et al. (2025) also detected multiple copies of the nuclear gene *LFY* (Figure 2A) which confirms the allopolyploid origin for *T. semaphora* as first suggested in Dawson et al. (2007). Based on the data currently available, it is impossible to determine which species were involved in the formation of *T. semaphora*. However, since the closest sequenced species on the maternal side is the Australian endemic *T. mucida*, (Jones et al. 2025 and Figure 2B), it is likely that *T. semaphora* originated in Australia and subsequently dispersed to NZ. This origin was earlier hypothesised for *T. semaphora* by Molloy & Dawson (1998) when its chromosome numbers were first reported. The affinity between *T. semaphora* and its Australian and NZ congeners is illustrated in Figure 2A & B.

#### Selected specimens studied

NORTH ISLAND: Aupouri, Sweetwater Rd, near Kaitaia, *Molloy B. P. J. s*. *n*., 03 Nov 1988, CHR 636773. Aupouri, *ex*. Sweetwater Rd, near Kaitaia, [Cultivated at Lincoln, Landcare Research], *Molloy B. P. J. s*. *n*., 22 Nov 1988, CHR 518036. Aupouri, Sand Hills Rd, Ahipara, *Molloy B. P. J. s*. *n*., 18 Nov 1999, CHR 640956. Aupouri, Sandhills Road, Ahipara to Awanui, *de Lange P. J. PJ 529 & Crowcroft G. M.*, CHR 473041. Aupouri, between Kaitaia and Ahipara, Sandhills Road, *Molloy B. P. J. s*. *n*. 10 Oct 1989, CHR 648831. Aupouri, Sandhills Road, Ahipara, *Molloy B. P. J. s*. *n*. 18 Nov 1999, CHR 640955. Aupouri, *ex*. Ahipara Gumfields Historic Reserve, near Kaitaia, [Cultivated at Lincoln, Landcare Research], *Molloy B. P. J. s*. *n.*, 8 Nov 1990, CHR 584261 B [CHR 584261 A in spirit collection not seen]. Eastern Northland, McLeod Bay, W side of Whangarei Heads Road, *Forester L*. *J*. *s*. *n*. 26 Oct 2012. Auckland, between Hatfields Beach and Waiwera, *Young M*. *E*. *s*. *n*. 01 Nov 2008, AK 303848. Auckland, Waitakere, Cornwallis, N of Spragg Monument, *Gardener R*. *O*. *10390* & *de Lange P*. *J*., 27 Nov 2002, AK 340233.

#### Selected observations reported on iNaturalist

http://naturewatch.org.nz/observations/4431686,

https://www.inaturalist.org/observations/17373387,

https://www.inaturalist.org/observations/17749005,

https://www.inaturalist.org/observations/65074018,

https://www.inaturalist.org/observations/140351726, https://www.inaturalist.org/observations/187600556.

## Acknowledgements

This work was funded in part by an Hansjörg Eichler grant to HRJ, a grant from the APSF (19047) to CAL, and the New Zealand Native Orchid Group. HRJ also thanks the Te Papa Foundation for a MSc scholarship, and Ian Bradshaw for further supporting his studies. The authors thank everyone who helped with collecting samples for this study and provided expertise on *Thelymitra*: the NZNOG, NOS, Bill Campbell, Matt Ward, Indy Marshal, Marly Ford, Kevin Matthews, Andrew Townsend, Ian St George, and Peter de Lange. A special thanks to Ngāitakoto and George and Kaio Hooper for supporting our research and collecting specimens for us. We also thank curators and sta at the AK, CANB, CHR, MPN, and WELT herbaria for their help with loans, granting sampling permission, and the Kew’s DNA and Tissue Bank for sharing DNA samples of Australian *Thelymitra*.

